# Boronic acid derivative inhibits LexA mediated SOS response in Mycobacteria

**DOI:** 10.1101/2023.11.15.567215

**Authors:** Chitral Chatterjee, Gokul Raj Mohan, V.C Hariharan, Bhumika Biswas, Vidya Sundaram, Ashutosh Srivastava, Saravanan Matheshwaran

## Abstract

Antimicrobial resistance (AMR) properties of several pathogens pose as one of the major global concerns demanding continuous innovation in order to be controlled. The bacterial “SOS” response, regulated by LexA and RecA, contributes to AMR through advantageous mutations. Therefore, targeting LexA/RecA system with a novel inhibitor could suppress “SOS” response and may reduce AMR. However, RecA presents a challenge as a therapeutic target due to its conservation across species, including humans. Concurrently, LexA, being absent in eukaryotes, can be potentially targeted, in part, due to its association with “SOS” response and developing AMR. Our studies combining bioinformatic, biochemical, biophysical, and cell-based assays present a unique inhibitor of mycobacterial “SOS” response wherein we show that the inhibitor interacts directly with the catalytic site residues of LexA of *Mycobacterium tuberculosis (*Mtb), consequently hindering its cleavage, resulting in suppression of “SOS” response. We observed important genes under the “SOS” regulon to be down-regulated in the presence of the inhibitor along with a consequent reduction in the rate of mutation frequency of mycobacterial cells. In essence, this study may facilitate further research on potential LexA inhibitors, reducing mutational rates and offering promise in mitigating AMR.

## Introduction

The silent pandemic of antibiotic resistance is on the rise. Nearly 1.27 million people succumb to antibiotic resistance-related deaths every year [1]. Out of the 7.7 million deaths from bacterial infections, 1.6 million results from tuberculosis (TB) alone, caused by the bacteria, *Mycobacterium tuberculosis* (Mtb). This is a major cause of concern as drug-resistant TB cases increased by 3% within a span of one year (2020-2021) [2]. There is a 13% increase in total TB cases and 32% increase in multidrug resistant/rifampicin-resistant TB cases in India, annually [3]. Without better treatment and control strategies, insurmountable losses would be incurred in the future. The emergence of extensive multidrug-resistant strains has complicated the prospects of controlling and eliminating TB [4]. Therefore, an effective strategy could be to target the proteins involved in regulating the pathways that mediate mutagenesis by developing anti-mutagenic molecules [5].

The concerted action of DNA damage repair pathways and “SOS” response accounts for the increased mutability, adaptability, and emergence of drug resistant pathogens [6-9]. Over the last decade, efforts aimed at targeting the “SOS” response have been gathering momentum to strengthen therapeutic efficacy [10-12]. Blocking the “SOS” response impairs the ability of the bacteria to repair its damaged DNA in response to stress. Moreover, error-prone mutagenesis in response to “SOS” activation, which further leads to the development of drug-resistant mutations, can be avoided by directly inhibiting the activation of this pathway [13, 14]. Screening and characterization of “SOS” inhibitors can help us in targeting the stress adaptation-DNA repair-mutagenesis axes of Mtb [15]. This can help in augmenting the already existing drug regimen and act as an adjuvant therapy to counter multidrug resistance.

“SOS” response in bacteria is regulated by two master regulator proteins - LexA (transcriptional repressor, which upon damage undergoes autoproteolysis to activate the “SOS” pathway) and RecA (activator, which upon DNA damage catalyzes the autoproteolysis of LexA) [16-22]. Inactivating the master-regulator proteins controlling this pathway, namely the RecA*/LexA axis has been shown to cause decreased mutagenesis post-antibiotic treatment, decreased minimum inhibitory concentration (MIC) of DNA damaging antibiotics, and re-sensitization of drug-resistant strains [8, 23, 24]. Inhibitors of RecA from both *E. coli* and Mtb have been identified [7, 25]. For example, Suramin has been demonstrated to be a potent inhibitor of bacterial RecA proteins, augmenting the antimicrobial properties of ciprofloxacin [7]. RecA inhibition in bacteria depleted of DNA gyrase results in a reversion of persistence and enhanced efficacy of antibiotics [26, 27]. Homologs of RecA are known to exist in most prokaryotic and eukaryotic organisms which makes targeting it challenging. Recently, efforts are being undertaken on targeting the other master regulator, LexA, which is absent in eukaryotes [8, 28, 29]. A test of the serine protease model). An extensive collaborative study resulted in the identification of novel inhibitors targeting *E. coli* LexA autoproteolysis [8]. Since LexA plays a crucial role in the “SOS” response of Mtb, inhibitors of Mtb LexA would directly target the mutagenesis and drug resistance axes. Recently, 3-aminophenylboronic acid (3-aPBA) has been reported to inhibit the autoproteolysis of *E. coli* LexA, which is known to differ from Mtb LexA [28]. The kinetics of interaction between *E. coli* LexA and 3-aPBA had not been determined in this study. The anti-mutagenic potential of the molecule had not been studied in detail. Further insights into the mechanistic aspect needed probing. Presently, no known inhibitors of Mtb LexA have been identified and Mtb LexA differs from its *E. coli* counterpart in several ways [30]. Mtb LexA harbours additional stretches of amino acids at its N-terminal end and linker region [31]. Unlike its *E. coli* counterpart, Mtb LexA binds to different “SOS” box sequences with comparable nanomolar affinity *in-vitro* [30]. Since boronic acid class of inhibitors have been reported to block the catalytic activity of serine protease family of proteins including Mtb LexA, we hypothesized that this class of inhibitors could exert a similar effect on Mtb LexA [16,32]. Through computational analysis, biochemical, biophysical, and cell-based assays, we identified a potential inhibitor of Mtb LexA which was found to be effective in preventing its autoproteolytic cleavage, resulting in stalling of the “SOS” response. Further, we also observed a significant decrease in the mutation frequency of mycobacterial cells and down-regulation of important genes under “SOS” regulon. This study reveals the inhibition of mycobacterial “SOS” pathway by a novel potential inhibitor. Essentially, it holds promise as an anti-mutagenic agent which may strengthen the current arsenal to boost anti-TB therapeutic strategies.

## Results and Discussion

### Screening of compounds to identify a potential inhibitor of mycobacterial “SOS” response

To identify a potential Mtb LexA inhibitor, first we performed molecular docking and then molecular dynamics (MD) simulations. Considering that boronic acid-based compounds have previously demonstrated inhibition of *E. coli* LexA [28], we evaluated several compounds for their binding to Mtb LexA, including three FDA approved drugs (**Table S1**). The rationale for selecting these compounds was that the boron moiety can form a covalent bond with the hydroxyl group of a serine residue in the active site of serine hydrolases. We predicted that this mechanism might apply to LexA, a known serine protease [32,33]. Therefore, we performed non-covalent docking of these compounds at the active site of Mtb LexA (see **Methods**).

All compounds exhibited low docking scores, indicating weak binding (**Table S1**). To further test the stability of these compounds in the active site, we performed molecular dynamics simulations, which revealed the instability of these compounds leading to their exit from the binding pocket (**Figure S1a**). All compounds exhibited low docking scores, indicating weak binding (**Table S1**).

To further test the stability of these compounds in the active site, we performed molecular dynamics simulations, which revealed the instability of these compounds leading to their exit from the binding pocket (**Figure S1a**).

Next, we employed covalent docking approach and assessed the ability of the compound to remain within the binding pocket after covalent interaction with a catalytic residue (S160). The compounds displayed varying covalent binding affinities and MMGBSA score (**Table S1**). We focused our analysis on 3-nitrophenylboronic acid (3-nPBA), with the most negative MMGBSA score and negative covalent docking affinity, as well as 3-aPBA, a compound previously shown to inhibit *E. coli* LexA through covalent interaction with the catalytic site residue [28].

The catalytic site residues (S160, K197) involved in the autoproteolytic activity of Mtb LexA were found to accommodate 3-nPBA, indicating that it may act by preventing autoproteolytic activity of Mtb LexA, thereby disallowing the repressor from dissociating from the cognate “SOS” boxes of LexA regulated genes. As shown in **Figure 1**, the boronate oxygen atoms of 3-nPBA and 3-aPBA were found to be involved in hydrogen bonds with the carbonyl oxygen of I157. Moreover, the π ring electrons of the phenyl ring of 3-aPBA interacts *via* cation-π interactions with K197 and boron makes ionic interaction with K197 in 3-nPBA (**Figure 1a, b**). The above-mentioned interactions with boronate oxygen atoms and phenyl rings of boronic acid derivatives have been observed previously [34].

**Figure 1.**
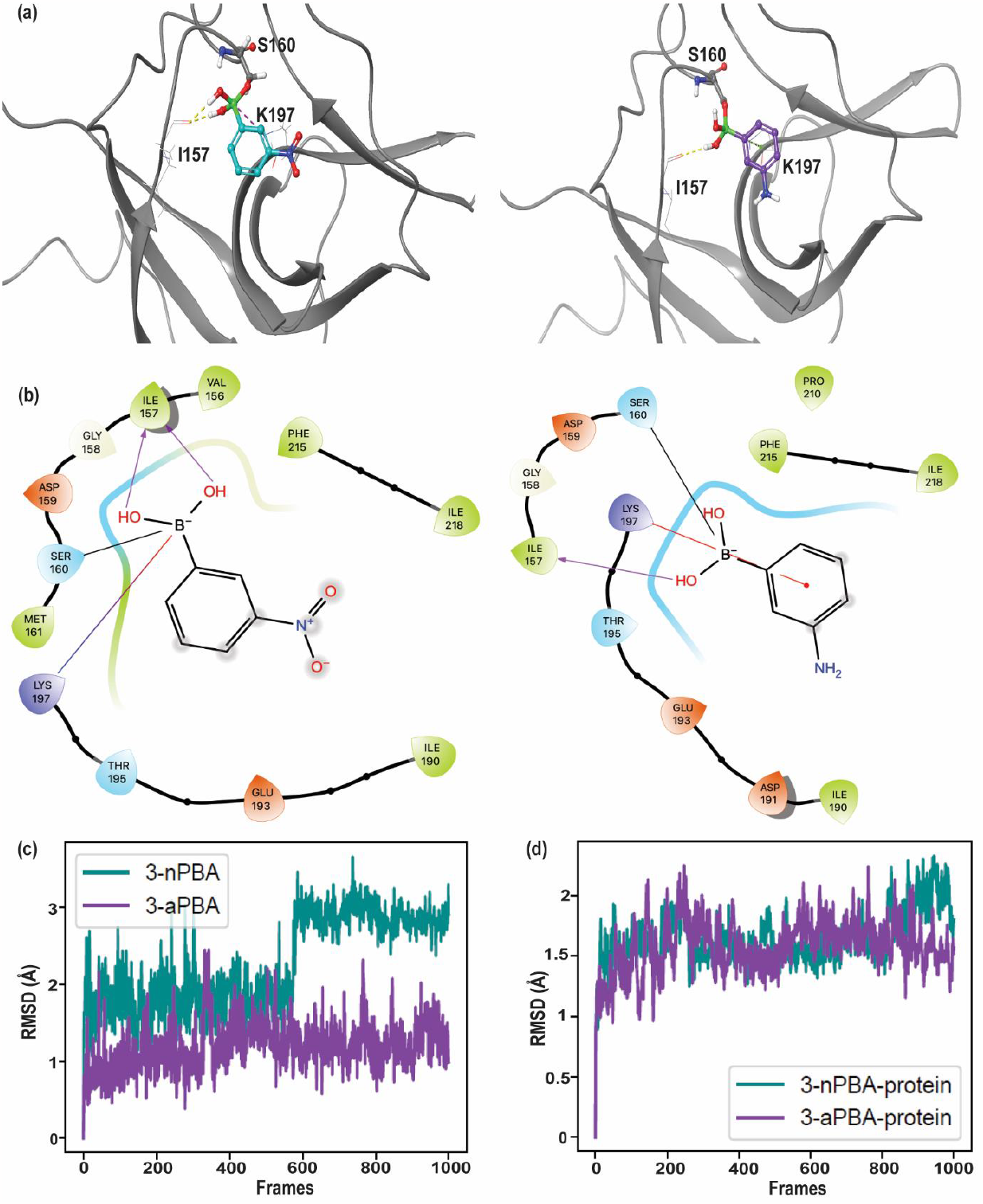
Interactions and stability of covalently docked 3-nPBA and 3-aPBA. (a) Docked pose of covalently bonded 3-nPBA (left) and 3-aPBA (right) in the binding pocket of Mtb LexA. Protein is shown in gray cartoon representation, while the ligands are shown in ball and stick representation. The interacting residues are represented in gray lines, reactive residue S160 in gray ball and stick representation. Hydrogen bonds and electrostatic interactions are shown as red and yellow dotted lines, respectively. (b) 2D ligand interaction diagram of 3-nPBA (left) and 3-aPBA (right). Hydrogen bonds are shown in purple arrows, electrostatic interaction is shown in blue-red shaded line and cation-pi interaction is shown as red line. (c) Root Mean Square Deviation (RMSD) of 3-nPBA and 3-aPBA during 10 ns (1000 frames) run. (d) RMSD of protein Cα atoms of 3-nPBA and 3-aPBA docked complexes during 10 ns (1000 frames) run.

In order to understand the stability of these covalently bound compounds, we performed MD simulations of these complexes. Both the compounds showed stable binding in the pocket over the course of simulations (**Figure 1c, d**). The interactions of 3-nPBA and 3-aPBA during the simulations showed very similar interactions. Most importantly, the compounds show ionic interactions with the catalytic K197 residue (**Figure S1**). This further suggests the mechanism of inhibition of LexA through these compounds. We then examined the binding energies of the two compounds during the simulations. 3-nPBA showed a more negative binding energy (-29.9 ± 4.1 kcal/mol) as compared to 3-aPBA (-23.2 ± 9.1 kcal/mol). This suggests that 3-nPBA might be a more potent inhibitor of Mtb LexA.

To experimentally validate these studies, we checked for the direct interaction of the protein with the inhibitor using isothermal titration calorimetry (ITC) and observed that 3-nPBA interacts with Mtb LexA with an affinity of 0.35 ± 0.26 mM (**Figure 2a, b)**. Moreover, to probe whether the catalytic site residues of the protein are involved in inhibitor binding as predicted by docking, we generated mutant(s) of the catalytic site residues using site-directed mutagenesis (SDM), and assessed their effect on inhibitor binding using ITC. Upon mutating both the catalytic site residues simultaneously (S160A and K197A), we found 10 times reduction in affinity for 3-nPBA (**Figure 2b**). This strongly suggests that mutating the residues responsible for autoproteolytic cleavage of Mtb LexA results in impaired inhibitor binding activity (**Figure S2)**. Experimental validation using ITC provided a strong indication that the inhibitor could directly interact with Mtb LexA to affect its function, a consequence of which may result in “SOS” inhibition.

**Figure 2.**
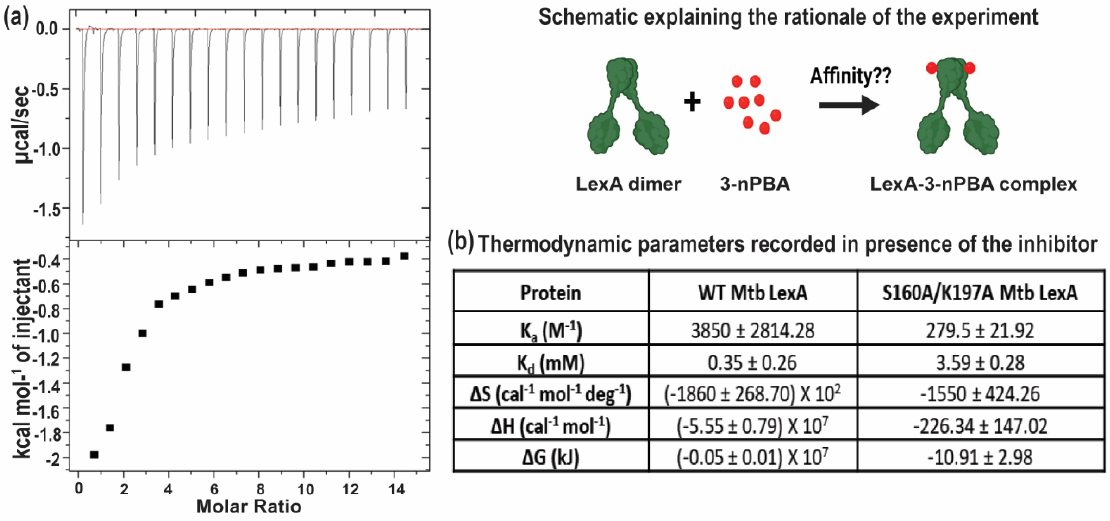
ITC analysis of Mtb LexA-inhibitor interaction. (a) Binding isotherm of WT Mtb LexA with 3-nPBA. Schematic explaining the rationale behind determining the kinetic parameters of interaction between Mtb LexA and 3-nPBA. (b) Thermodynamic parameters recorded in presence of 3-nPBA.

### Biophysical and biochemical studies reveal that 3-nPBA stabilizes Mtb LexA

To gain more insight into the changes in secondary and tertiary structure of Mtb LexA in presence of the inhibitor, we performed circular dichroism and extrinsic fluorescence-based studies wherein, we compared near UV spectra of Mtb LexA alone and in presence of 3-nPBA. Treatment with the inhibitor resulted in increase in alpha helical content of the protein as deduced from the appearance of a more prominent peak at 222 nm of the inhibitor-protein complex as compared to the spectra of the native protein alone (**Figure 3a**). Moreover, an increase in negative ellipticity is indicative of enhanced stability of the protein in the presence of the inhibitor.

**Figure 3.**
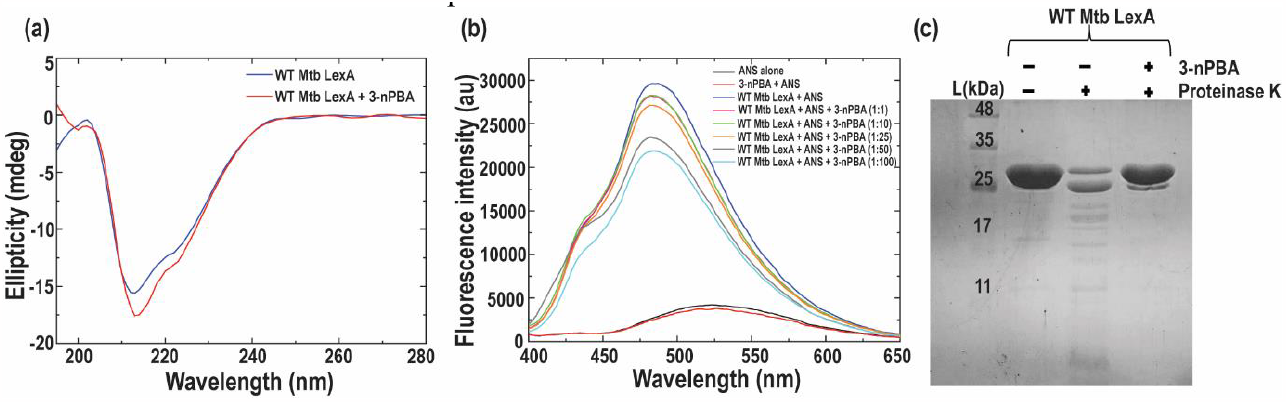
Characterization of Mtb LexA with potential inhibitor. (a) Secondary structural changes of Mtb LexA in presence of 3-nPBA as determined by CD Spectroscopy. (b) Tertiary structural changes of Mtb LexA in presence of increasing concentrations of 3-nPBA as determined by extrinsic fluorescence-based studies. Fluorescence intensity is shown in arbitrary units. (c) Proteinase K protection assay of Mtb LexA in the presence of 3-nPBA.

Extrinsic based fluorescence using ANS (8-Anilinonaphthalene-1-sulfonic acid) was performed and the obtained spectra of Mtb LexA in the presence of increasing concentrations of the inhibitor resulted in concentration dependent quenching in fluorescence (**Figure 3b**). As ANS is known to bind to the hydrophobic patches of the protein, elevated fluorescence intensity of ANS indicates increased unfolding of the protein to reveal the hydrophobic patches [35]. In this case, the protein possibly gets more stabilized in presence of the inhibitor, resulting in fluorescence quenching. We tested the stability of the protein in presence and absence of the inhibitor using a known protease like Proteinase K, and observed that the inhibitor protected Mtb LexA from undergoing cleavage (**Figure 3c**).

### 3-nPBA protects Mtb LexA from autoproteolysis without affecting dimerization

Next, we assessed the effect of 3-nPBA on the autoproteolytic activity of Mtb LexA. Corroborating with the previous observation, whereby we found Mtb LexA to get protected from Proteinase K mediated cleavage, we found the inhibitor to display a protective effect on the protein, preventing it from undergoing autoproteolysis as well (**Figure 4a**). LexA is known to undergo autoproteolysis at alkaline pH [16, 36]. Mutating the K197 residue alone prevented autoproteolysis. This was further observed for the K197A/S160A double mutant, wherein both residues directly involved in autoproteolysis were mutated (**Figure 4a**).

**Figure 4.**
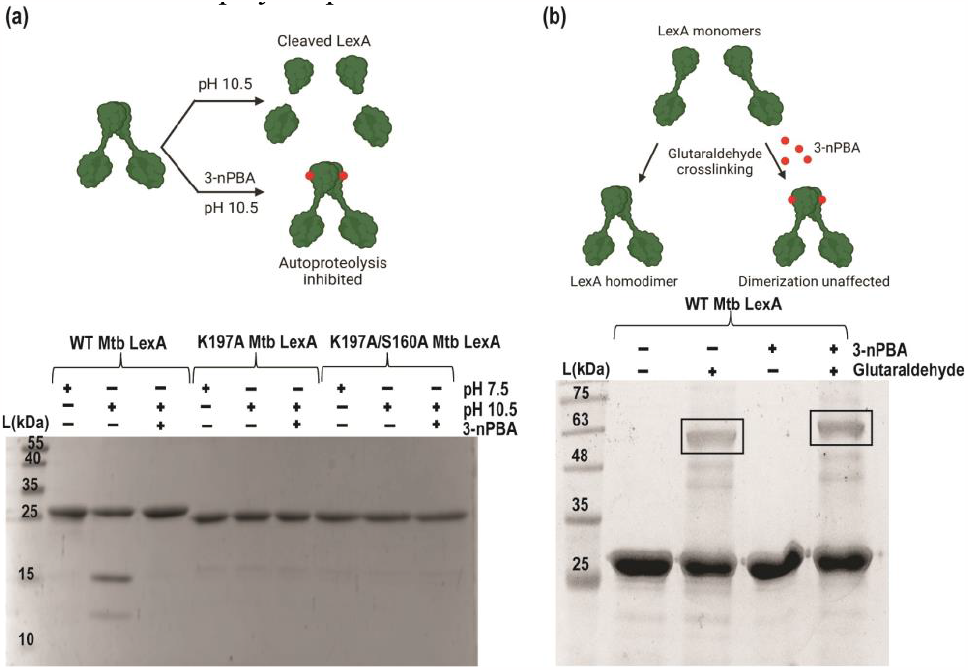
3-nPBA protects Mtb LexA from autoproteolysis without affecting dimerization. (a) Autoproteolytic cleavage assay of Mtb LexA and its mutants in presence of 3-nPBA. (b) Glutaraldehyde cross-linking assay of Mtb LexA in presence and absence of “SOS” inhibitor, 3-nPBA reveals no change in dimerization state of the protein.

We hypothesized that since the inhibitor could prevent cleavage of Mtb LexA and stall “SOS” response, it may also be playing an active role in stabilizing the LexA-DNA complex. LexA has been reported to bind as a dimer to its cognate “SOS” boxes with nanomolar affinity [30, 37]. Hence, we examined whether the inhibitor affected this dimerization property of Mtb LexA and observed through cross-linking assays that dimerization of the protein remained unaffected in the presence of the inhibitor (**Figure 4b)**.

### “SOS” inhibitor stabilizes Mtb LexA-DNA complex

Real-time kinetic studies were performed using biolayer interferometry (BLI) to assess how the inhibitor would affect Mtb LexA-DNA interaction. The DNA used here contained the “SOS” box of *dnaE2. dnaE2* gets expressed upon induction of “SOS” response and is implicated in error-prone mutagenesis [38-40]. Although the association constant (k_on_) of the Mtb LexA-DNA interaction did not alter substantially when compared between the untreated and inhibitor-treated conditions, there was a significant reduction by nearly 23 times in the dissociation constant (k_off_) in case of the latter (**Figure 5a, c**). We observed that in the case of the wild-type protein-DNA interaction, there was an increase in DNA binding affinity by nearly 16 times in the presence of 3-nPBA when compared to the untreated condition. Contrarily, in the mutant protein, this difference in the dissociation rate was not significant (**Figure 5b, c**). Finally, the cumulative affinity of the mutant towards DNA decreased in the presence of 3-nPBA by 1.5 times when compared to the untreated condition.

**Figure 5.**
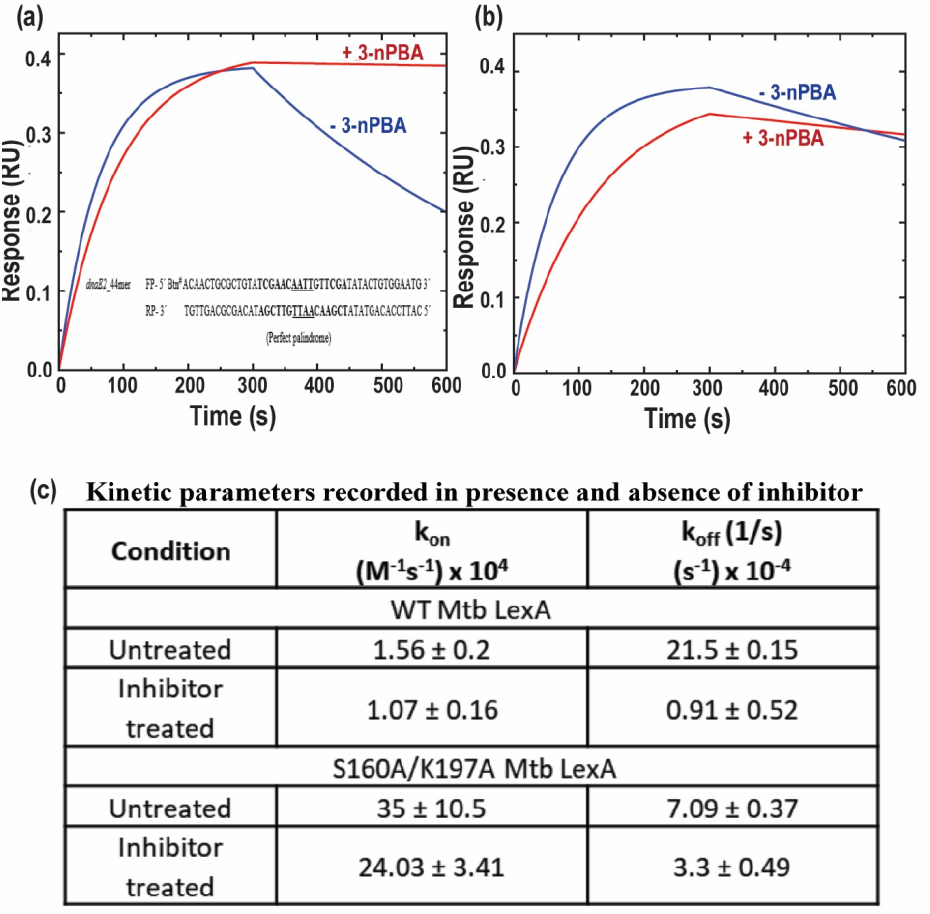
Dissociation of Mtb LexA assessed in presence of “SOS” inhibitor, 3-nPBA. (a) Slower dissociation of Mtb LexA from DNA seen in the presence of inhibitor as revealed by real-time kinetic studies. (b) No significant change in the dissociation rate of S160/K197A Mtb LexA from DNA was seen in the presence of the inhibitor compared to that observed in (b). (c) Table showing kinetic parameters recorded for the interactions in (a) and (b).

We observed that the dissociation rate of Mtb LexA from its target DNA sequence decreased in the presence of the inhibitor, demonstrating the stabilization of Mtb LexA-DNA interaction in the presence of 3-nPBA (**Figure 5a, c)**. To summarize, Mtb LexA binds to 3-nPBA with its catalytic site residues, which prevents its cleavage and also stabilizes its association with DNA. This consequently would affect the repressor protein from falling off the “SOS” boxes, thereby maintaining the suppression of “SOS” responsive genes.

### Inhibitor mediated suppression of mycobacterial “SOS” response

We generated a fluorescence-based “SOS’’ reporter construct containing the promoter region inclusive of a consensus “SOS” box sequence of *dnaE2* to visualize the “SOS” response in mycobacteria. We hypothesized that mycobacterial LexA would bind to the “SOS” box under normal conditions. Upon DNA damage and induction of “SOS” response, LexA undergoes autoproteolysis and falls off, thereby de-repressing the downstream reporter gene (in this case, *mCherry*) (**Figure 6a**). Using this “SOS’’ reporter construct, we assessed the “SOS’’ inhibitory activity of 3-nPBA. As expected, we found mycobacterial “SOS’’ activation to be compromised in its presence even at the sub-inhibitory concentration of the inhibitor **(Figure 6b)**.

**Figure 6.**
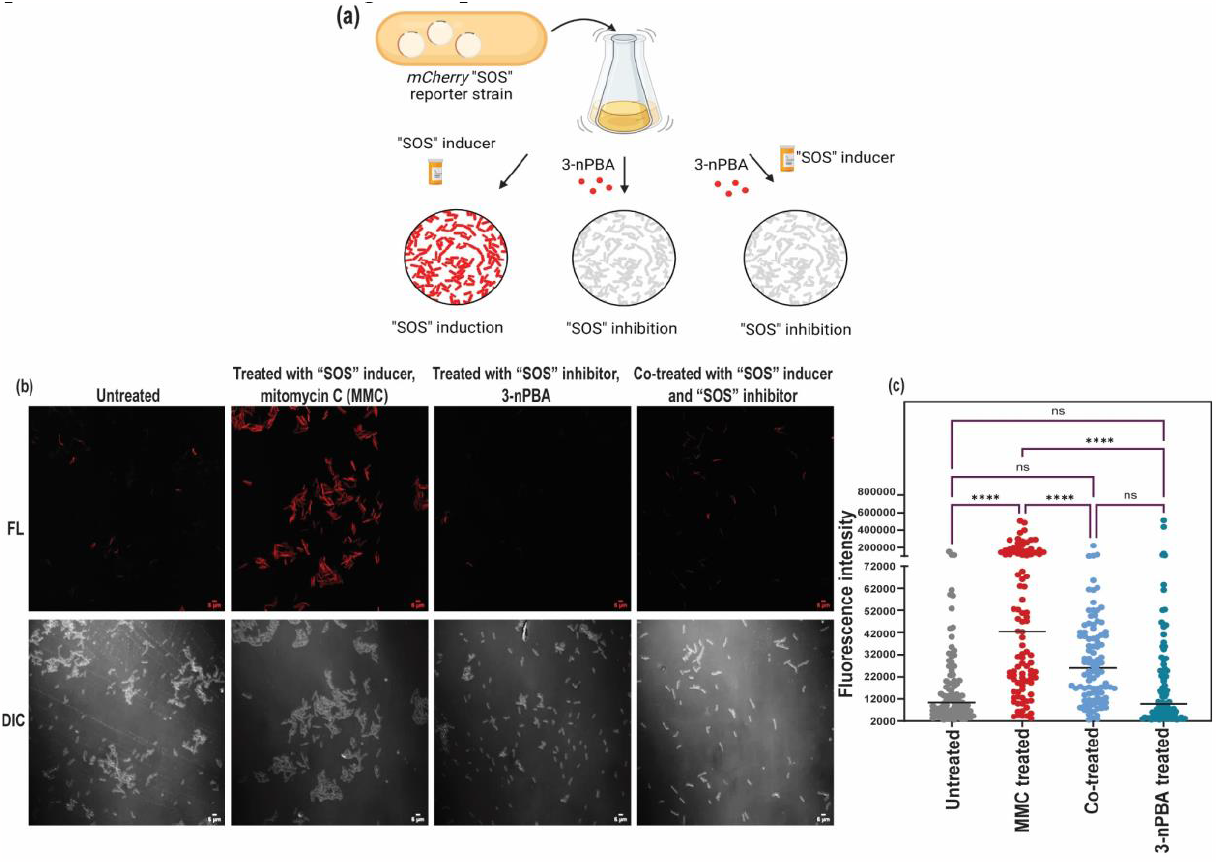
Assessment of “SOS” inhibitory activity of the potential inhibitor, 3-nPBA using fluorescence-based reporter constructs. (a) Strategy used to construct the fluorescence-based mycobacterial “SOS” inducible reporter. (b) Expression of *mCherry* in the presence of a known “SOS” inducer, mitomycin C can be observed in contrast to the cotreated cells with the potential “SOS” inhibitor, 3-nPBA, whereby “SOS” induction gets repressed as observed in the representative confocal microscope images. (c) Quantification of cells under varying treatments. One-way ANOVA was performed (****=p<0.0001, ns=non-significant).

Mitomycin C is a known inducer of “SOS” response and it functions by crosslinking DNA and inducing double strand breaks [41]. While “SOS” induced samples (treated with mitomycin C) harbored cells expressing *mCherry*, those treated with the inhibitor alone or when co-treated with both the inhibitor and inducer exhibited basal level fluorescence, indicative of sustained “SOS” inhibition. It is important to mention here that cells exhibit a basal level of “SOS” response and even in the untreated condition, we can observe some cells expressing *mCherry* **(Figure 6b)**. The cells that showed highly reduced fluorescence upon treatment with the inhibitor at sub-inhibitory concentrations were still viable as determined from the Resazurin Reduction Assays (REMA) (**Figure S4**). To further explore this, we used a constitutively expressing *mCherry* expressing reporter, treated the cells with the inhibitor at the same sub-inhibitory concentration as used during the microscopic analysis with the fluorescence-based “SOS’’ reporter and checked for fluorescence. The treated cells continued to exhibit fluorescence, indicative of their viability **(Figure S3)**. Hence, we can conclude that treatment with the inhibitor indeed suppressed “SOS” response without affecting cell survival. We also probed for the bactericidal activity of 3-nPBA against different bacterial strains-*M. smegmatis*, the avirulent strain and the virulent strain of Mtb (Mtb H37Ra and Mtb H37Rv, respectively), and against representative Gram-positive and negative organisms **(Figure S4)**. Moreover, cytotoxicity assessment studies conducted on macrophage cell line RAW264.7 revealed that even at 29X of the MIC of the inhibitor, no observable cytotoxic effect could be observed **(Figure S5)**. Hence, 3-nPBA can be considered non-cytotoxic for mammalian cells even at higher concentrations.

### 3-nPBA curbs expression of “SOS” regulon genes and is anti-mutagenic

To check the efficacy of 3-nPBA on the mutation frequency of antibiotic treated cultures, we performed mutation frequency tests. Ciprofloxacin was taken as a positive control. Ciprofloxacin is a well-known “SOS” inducer which results in increased expression of DNA damage inducible genes forming a part of the “SOS” regulon whose concerted action brings about mutagenesis [42]. Expression levels of the “SOS” inducible gene *dnaE2*, a primary contributor of bacterial error-prone mutagenesis, remained high in ciprofloxacin induced persisters even after 30 hours of culturing in antibiotic-free medium [39]. To test whether 3-nPBA acts as an anti-mutagenic molecule, we subjected wild-type *M. smegmatis* to treatment with ciprofloxacin alone, 3-nPBA alone, ciprofloxacin in combination with 3-nPBA with the untreated negative control. Next, we recovered the cultures in antibiotic free medium while reviving the co-treated cultures in the presence of the inhibitor and subsequently plating them all on ciprofloxacin-containing plates to assess the mutation frequency rate. We found that mutation frequency decreased by 14 times in 3-nPBA co-treated samples as compared to the untreated control. Treating with ciprofloxacin alone resulted in 3 times higher mutation frequency compared to the untreated control (**Figure 7a**). We also attempted to delineate the underlying mechanism by assessing the differential expression of selected “SOS” regulon genes upon treatment with 3-nPBA. As expected, treatment with the inhibitor resulted in the down-regulation of the genes that are highly controlled by LexA, implying that even at the level of transcription, the inhibitor remains effective in stalling mycobacterial “SOS” response (**Figure 7b**).

**Figure 7.**
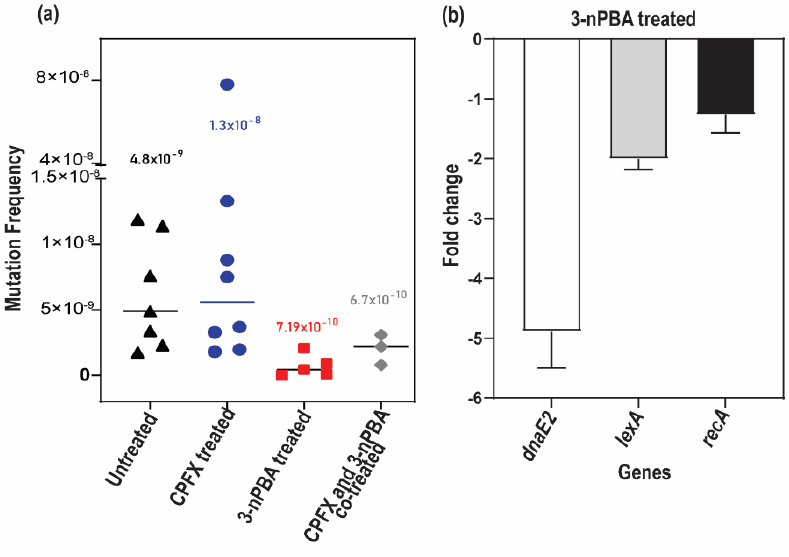
Deciphering the anti-mutagenic effect of “SOS” inhibitor 3-nPBA from differential gene expression studies and mutation frequency analysis. (a) Comparative analysis of mutation frequency on ciprofloxacin (CPFX) containing plates upon treatment of cells in presence of “SOS” inducer (ciprofloxacin), “SOS” inhibitor (3-nPBA), and in presence of both ciprofloxacin and “SOS” inhibitor, 3-nPBA with respect to that of the untreated mycobacterial cells. 9 biological replicates were taken for each condition and values were plotted using GraphPad Prism. (b) Down-regulation of mycobacterial “SOS” regulon genes upon treatment with 3-nPBA using qRT-PCR. The results shown are from 3 biological replicates, each with 3 technical replicates.

## Conclusion

There is an overwhelming need to formulate new strategies to counter mycobacterial multidrug resistance [43]. The current antibiotic arsenal needs strengthening by complementing with anti-evolution molecules which can interfere with the bacteria’s ability to acquire drug resistance [44]. Since the bacterial “SOS” response pathway is one of the major drivers of its drug resistance, [45-47] targeting the master-regulators controlling this pathway can prove to be promising and scientists have started conducting their research in this direction. The identification of small molecules which can inhibit the “SOS” response activation have opened up new avenues in the field of antimicrobial therapy [5]. The suppression of mycobacterial “SOS” response by targeting one of the master-regulators, RecA, has been found to slow drug resistance [48]. However, since RecA bears mammalian homologs, targeting the other master-regulator, LexA deemed to be more attractive. Currently very few inhibitors have been identified to target the LexA/RecA axis [49] and the unavailability of Mtb LexA inhibitors prompted us to execute this study.

We conducted our search for an effective inhibitor of Mtb LexA, the repressor master-regulator controlling “SOS” response. Since boronic acid derivatives were established as prominent inhibitors of LexA autoproteolysis, we hypothesized that modifying the side-chains of the boron-containing inhibitors could lead to the identification of efficacious Mtb LexA inhibitors. Through our study, we characterized a potential boronic acid-containing Mtb LexA inhibitor that proved effective in stalling “SOS” induced mutagenesis in mycobacteria.

Cell-based screening assays using damage-inducible reporter strain was tested in the presence of known mutagen and potential anti-mutagenic inhibitor molecule to check for their activity. Since 3-nPBA was found to be effective in stalling “SOS” and had a more potent killing activity than another derivative, we chose to proceed with it for subsequent studies. 3-nPBA was found to be potent against multiple bacteria, which establishes its sufficient breadth across species role in stalling “SOS” response. Moreover, its non-cytotoxic nature also proved to be encouraging for designing further studies.

We found that the mutation frequency decreased by 14 times in 3-nPBA co-treated samples as compared to the untreated control. Treating with ciprofloxacin alone resulted in 3 times increase in the mutation frequency compared to the untreated control. These results indicate that the “SOS” inhibitor is indeed anti-mutagenic. Studying the gene expression patterns of mycobacterial “SOS” regulon genes in the presence of the inhibitor further validated our findings as they were found to be down-regulated in its presence, thereby suppressing “SOS” mediated mutagenesis.

Through covalent docking studies, we found the catalytic side residues (S160 and K197) of Mtb LexA to be possibly involved in interacting with 3-nPBA. We generated the catalytic mutant(s) of Mtb LexA, assessed the binding to 3-nPBA and compared it with that of the wild-type protein. The binding studies using ITC revealed compromised inhibitor binding properties of the double mutant. The covalent bond formed between the inhibitor and the protein ensures the irreversibility of the interaction and this satisfies the criteria to make the cut for a promising inhibitor molecule [50].

Further, we performed biochemical and biophysical studies to elucidate the effect of the inhibitor on Mtb LexA. Proteinase K assays to compare the stability of the protein in presence and absence of the inhibitor revealed that 3-nPBA has a stabilizing effect on Mtb LexA. There was no effect though on the dimerizing ability of Mtb LexA in its presence. We observed changes in the secondary structure of the protein in the presence of the inhibitor using secondary structure analyses.

Concentration-dependent change in tertiary structure of the protein in the presence of increasing concentrations of the inhibitor was observed using ANS-based extrinsic fluorescence assays. We found that in the presence of 3-nPBA, the dissociation of Mtb LexA from DNA slowed using real-time kinetic studies. All these studies taken together provide us a detailed understanding of how this first-of-its-kind mycobacterial “SOS” inhibitor may help preventing the bacteria from gaining AMR.

To date, we did not have anti-mutagenic inhibitors that could target the “SOS” response axis of mycobacteria. Such inhibitor molecules hold promise in adjuvant therapy to accentuate the activity of existing drug regimens. This study lays the platform for developing an anti-mutagenic “SOS” inhibitor screening platform that may help target not just *Mycobacterium tuberculosis*, but also other pathogenic Gram-positive and Gram-negative bacteria, which is much needed in this era of expanding antimicrobial resistance.

## Methods

### Reagents, plasmids and strains

Bacterial plasmids and strains used in this study are mentioned in **Table S2 of Supporting Information**. Primer sequences are listed along with the constructs generated in the study in **Table S3 of Supporting Information**. All the chemicals, reagents, and media, were purchased from Hi-Media, Sigma Aldrich, SRL, and Difco, and enzymes were purchased from New England Biolabs, Genei and Promega. 50 mg/ml stocks of the inhibitors (3-nPBA and 3-aPBA) were prepared in 40% DMSO.

### Growth conditions

*M. smegmatis* was grown in 7H9 media with 0.2 % Tween-80 (v/v), 0.5 % glycerol. For growing Mtb H37Ra and Mtb H37Rv, OADC was used as a supplement in 7H9. 7H11 agar with 0.5 % glycerol was added for solid media. The concentrations of antibiotics used for mycobacteria were 25 μg/ml kanamycin and 50 μg/ml hygromycin as and when required. Strains were grown at 37°C. LB was used for culturing *E. coli* and *S. aureus* and while performing resazurin reduction assays, secondary cultures were grown in MH broth.

### Confocal microscopy

Mycobacterial cells bearing the “SOS” inducible reporter were grown upto O.D_600_ 0.4 and divided into different tubes containing 40 ng/ml of “SOS” inducing agent such as mitomycin C or inhibitor, 3-nPBA at one-fourth of its MIC or in combination of both the “SOS” inducer and inhibitor and grown for 4 h before analysis. Cells were washed with 1X PBS, fixed with 3 % paraformaldehyde (PFA), and mounted on agar padding before imaging. Samples were observed under 63X magnification of the confocal microscope. Samples were excited at 565 nm and an emission filter of 610 nm was taken for observing *mCherry*-expressing cells.

### Mutation frequency analysis

Protocol followed as per Salini et al., 2022 [39]. Additionally, modifications suitable for the experiment were as follows: Concentration of 3-nPBA tested: ½ MIC concentration against *M. smegmatis* ie, 15 μg/ml (89.8 μM). Concentration of ciprofloxacin tested: 7X less than the molar concentration of that of the inhibitor used ie, 12.8 μM. Incubation was done for 6 days post which plating was carried out. For viability testing, 10^5^ and 10^7^ dilutions of saturated cultures were plated while the remaining culture was plated as mentioned in the protocol on antibiotic containing plates.

### RNA isolation, DNase treatment, cDNA conversion and qRT PCR

For this, standard procedure as given in [51] with few modifications was followed. As a modification to the mentioned protocol, introduced addition of 0.5 mm Zirconia beads after addition of Trizol and Chloroform. Samples were intermittently vortexed and kept on ice. Quick vortexing for 10 secs for each sample for a total of 3 mins with intermittent incubation on ice was done to ensure proper lysis. 25 ml of the mycobacterial cultures were taken for RNA isolation. DNase treatment using DNase I (Promega) was performed according to manufacturer’s instructions. 1 μg of RNA from each sample was converted to cDNA following instructions of Promega. qPCR was performed using SyBr green. 65°C was chosen as annealing temperature. *rpoB* was used as house-keeping control and reference.

### Three-dimensional structure of C-terminal domain (CTD) of Mtb LexA: selection and preparation

At the time of this study, only a single study on X-ray structure of the C-terminal domain (CTD) of Mtb LexA [36] had been done and the data corresponding to that has been deposited in Protein Data Bank (PDB). The crystal structure of the CTD has four forms; forms I (PDB ID - 6A2Q) and II (PDB ID - 6A2R) are both unmutated with one and six monomers in the asymmetric units, respectively. The crystal structure with PDB ID 6A2Q with the monomeric CTD was selected for further investigations. The CTD structure was generated using the Protein Prep wizard [52] the protein preparation step included the addition of missing residues and hydrogens, optimization of hydrogen bonds, removal of water molecules, and energy minimization with convergence to a maximum RMSD of 0.3 Å. Although the crystal structure of CTD with a co-crystallized inhibitor is yet to be determined, a probable binding pocket based on the comparative analysis of the CTD of *E. coli* and *P. aeruginosa* has been suggested [16]. The immediate neighborhood of the catalytic residues forms a pocket comprising of residue stretches 154-161 and 193-197 that could be considered as a target for the development of inhibitors; stretch 154-161 is substantially the same in LexA from Mtb,

*E. coli* and *P. aeruginosa* and the stretch 193–197 is highly conserved in *E. coli* and *P. aeruginosa*. Accordingly, the energy-minimized protein structure was then processed in the Glide module [53, 54] to generate grids with the centroid defined by residue stretches 154-161 and 193-197.

### Ligand Preparation

LigPrep with Epik [55,56] was employed to generate the three-dimensional structures of the ligands mentioned in **Table S1**. The generated structures were then optimized at pH 7.0 ± 0.5. The processed protein structures with the grid and the ligands were subjected to two docking procedures; conventional molecular docking in the Glide module and covalent docking using the CovDock module [57] of Schrodinger, both at extra-precision (XP) mode [54].

### Covalent Docking

For covalent docking involving boronic acid derivatives, the reaction type and reactive residue on the receptor were identified as boronic acid addition and the catalytic S160, respectively. Further, the CovDock module of Schrodinger was used to dock the molecules to the catalytic site of Mtb LexA. The CovDock affinity score and MMGBSA score was reported for all the molecules (**Table S1**).

### Molecular Dynamics Simulations

To analyze the stability of the protein-ligand complexes, molecular dynamics (MD) simulations of the selected complexes were performed using Desmond [58]. Each of the selected protein-ligand complexes was simulated in a truncated octahedron box solvated with explicit TIP3P water molecules. OPLS3e force field [59] was used to model the protein and inhibitor molecules. The default energy minimization and equilibration settings of Desmond were applied to the simulation system. The equilibrated systems were used for the final production run of 10 ns using the NPT ensemble at 310 K and 1 atm pressure. The stability of the protein-ligand complexes was analyzed using the “simulation integration diagram” and “simulation quality analysis” tools of Desmond.

### Over-expression proteins and purification of

Mtb LexA construct generated in our previous study was used [30]. Mutant was generated by the non-overlapping site-directed mutagenesis. The primers used have been listed in **Table S3 of Supporting Information**. All constructs generated were confirmed by sequencing. The recombinant WT and its mutant proteins were over-expressed in *E. coli* BL21(DE3) cells following the published protocol [30].

### Isothermal Titration Calorimetry (ITC)

The recombinant proteins were titrated against the ligand in Microcal ITC 200 (GE). 20 injections, each of 2 μl, were made at 150 s intervals, at 25°C. The heat of the reaction per injection (microcalories per second) was determined by the integration of the peak areas. The concentration of protein has been calculated taking into consideration its dimeric form in solution. 25 mM phosphate buffer, pH 7.5 was used.

### Circular dichroism

Circular dichroism (CD) studies were performed using Jasco J-815 spectropolarimeter according to our previously published protocol [30]. About 5 μM of protein in 10 mM HEPES, 50 mM NaCl (pH 7.5) and 100X molar concentration of the inhibitor was taken for analysis. The data shown are an average of three independent scans after correcting for the buffer baseline. Origin 8.1 software was used for plotting the recorded spectra.

### Extrinsic fluorescence

Extrinsic fluorescence-based studies were performed according to our previously published protocol [30]. About 5 μM of the protein in 10 mM HEPES, 50 mM NaCl (pH 7.5), was incubated with varying concentrations of the inhibitor (3-nPBA) for 30 mins at 37°C. All measurements were corrected for fluorescence intensity of buffer, inhibitor, and ANS intrinsic fluorescence.

### Proteinase-K protection assay

Proteinase K cleavage of WT Mtb LexA was carried out in the presence of 3-nPBA. 1 mg/ml protein was pre-incubated for 15 mins with the inhibitor (1:100 ratio) in a buffer composed of 10 mM HEPES, 50 mM NaCl (pH 8), and 0.001 % proteinase K was used. The assay was carried out at 37°C. After 4 mins, 10 μl of the sample was taken, and proteinase K was inactivated with 1 mM PMSF and 5X SDS dye. The prepared samples were analyzed on a 15 % SDS-PAGE.

### Autoproteolysis cleavage assay

Autoproteolytic cleavage of Mtb LexA and its variants were induced using 100 mM CAPS, 300 mM NaCl (pH 10.5) and by incubating the proteins at 37°C for 6h. 5 μM of each protein was pre-incubated for 3h with the inhibitor (1:100 ratio) and then subjected to autoproteolysis. Samples were analyzed on a 15 % SDS-PAGE.

### Cross-linking reactions

The cross-linking reactions were performed according to already standardized protocol [30] either in presence or in absence of the inhibitor (3-nPBA). 1:100 was taken as the ratio of protein to inhibitor. After stopping the reactions with 25 mM of DTT, samples were prepared and separated on a 12 % SDS-PAGE.

### Biolayer interferometry (BLI)

The interaction studies between Mtb LexA with biotinylated ds 44mer of *dnaE2* “SOS” box (sequence listed in **Table S3 of Supporting Information**) in presence or absence of the inhibitor, 3-nPBA in a 1:10 ratio of protein: inhibitor was performed using biolayer interferometry according to previously published protocol [30]. The proteins were passed onto the chip and change in response units (RU) was analyzed both in presence and absence of the inhibitor. After reference data subtraction, 1:1 binding model was applied for fitting and plotting the data.

## Acknowledgments

The authors are grateful to members of SAIF, CDRI for permitting usage of the BLI Facility, Dr. Garima Khare for facilitating the use of BSL-III Facility at the University of South Campus, Delhi, the CD and Confocal facilities at IIT Kanpur. Authors thank Dr. Dharmaraja Allimuthu for providing synthesized 3-nPBA for preliminary studies. Authors thank Dr. Santosh K. Misra for permitting the use of his cell culture facility and Mr. Niranjan Chatterjee for assisting in the cytotoxicity assays. Some of the figures were prepared using BioRender. The authors thank Dr. Krishna Kurthkoti for the gift of pMV262∼*mCherry* and late Prof. M. Vijayan for the gift of S160A Mtb LexA construct. Authors thank Ms. Umang Gupta, Mr. Deepanshu Singla and Dr. Dharmaraja Allimuthu for helpful discussions during the study. Authors are also grateful to Dr. Appu K Singh and Dr. Soumitra Ghosh for critically reviewing the manuscript. SM is supported by DBT IYBA, SERB and STARS-MHRD, ICMR. CC acknowledges the Ministry of Human Resource Development, Government of India, for the fellowship. AS acknowledges support from DBT Ramalingaswami Fellowship, SERB Start-up Research Grant and IIT Gandhinagar for HPC facilities.

## Author contributions

CC and SM designed the experiments. CC performed the experiments. GRM and BB performed antibiotic susceptibility tests. HVC helped in confocal microscopy and mutation frequency tests. AS and VS performed all the docking studies and MD simulations. CC wrote the original draft. AS and SM edited the manuscript. SM supervised the study.

## Data availability statement

All supporting data and sequence information are included within the main article and its Supporting Information.

