## Supplementary materials for "Boronic acid derivative inhibits LexA mediated SOS response in Mycobacteria"

### Supporting information

**Table S1: Compounds screened in this study**

| Compound | Chemical structure | MMGBSA dG Bind | cdock affinity | Glide Score |
|---|---|---|---|---|
| Phenylboronic acid | 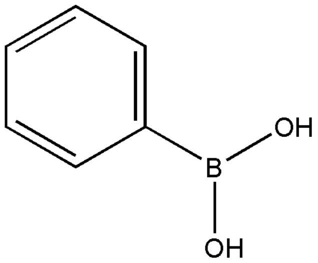 | -2.78 | -3.281 | -2.446 |
| 3-aminophenylboronic acid | 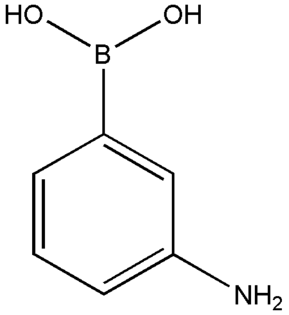 | -1.93 | -2.861 | -3.341 |
| **3-nitrophenylboronic acid** | 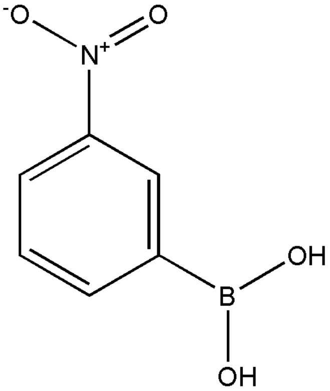 | **-4.96** | **-3.15** | **-3.267** |
| Benzene 1,4 di-boronic acid | 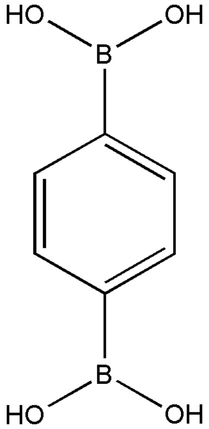 | -2.99 | -2.446 | -4.274 |

| | | | | |
|---|---|---|---|---|
| 4-pyridylboronic acid | 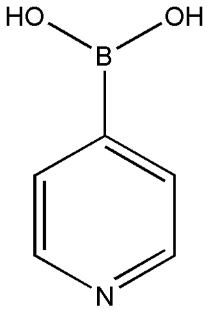 | 2.9 | -3.466 | -3.512 |
| 4-formylphenylboronic acid | 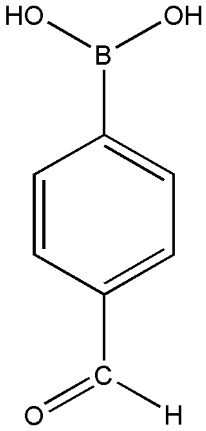 | 11.87 | -3.76 | -3.501 |
| 3-hydroxyphenylboronic acid | 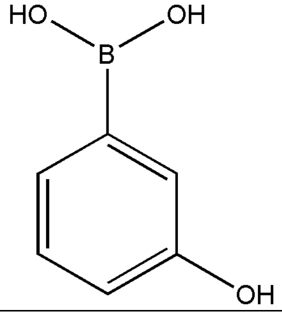 | 1.53 | -3.391 | -3.59 |
| 3-carboxyphenylboronic acid | 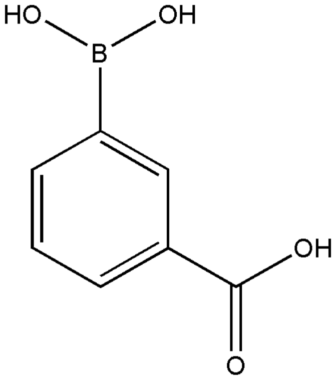 | 1.17 | -3.285 | -3.128 |
| [3-(2-carboxyvinyl)phenyl] boronic acid | 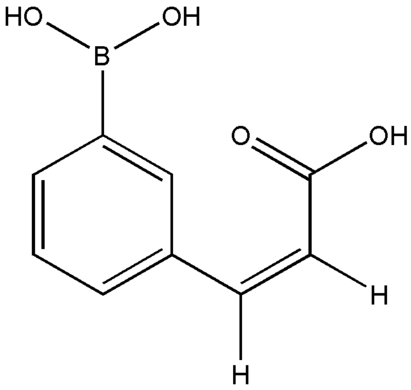 | 1.53 | -2.642 | -3.819 |

| | | | | |
|---|---|---|---|---|
| Vaborbactam | 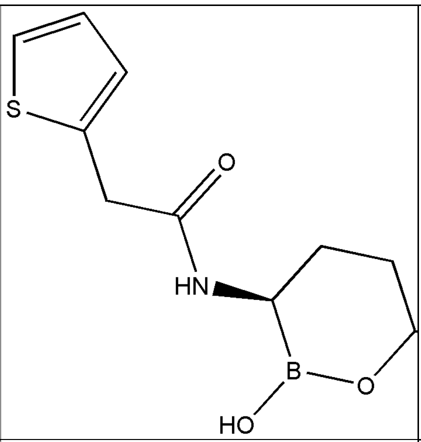 | Not determined | -3.713 | -3.463 |
| Bortezomib | 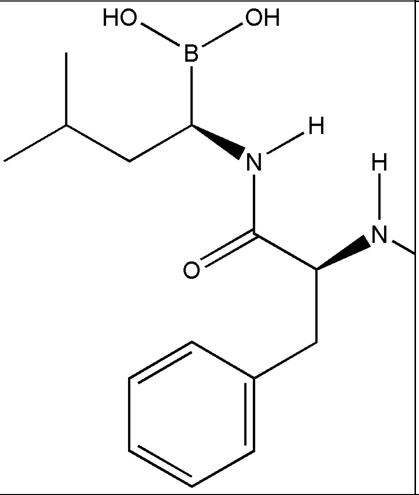 | 13.95 | -2.874 | -4.107 |
| Ixazomib | 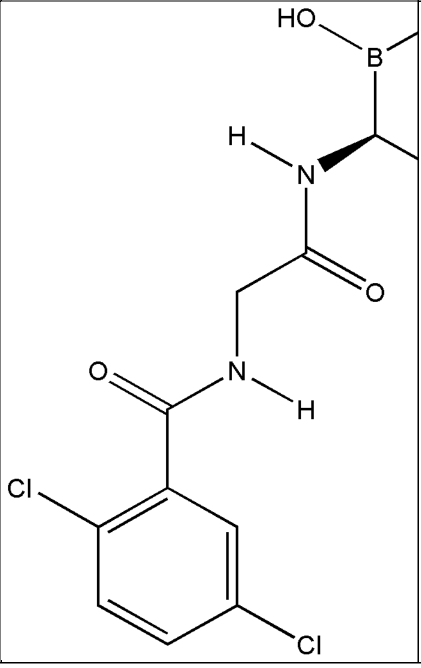 | 2.87 | -1.487 | -4.949 |

**Table S2: Bacterial strains and plasmids**

| Name | Characteristics | Source |
|---|---|---|
| **Bacterial Strains**<br>*E. coli* DH5α | Wild-type *E. coli* K12 strain. | Laboratory stock |
| *E. coli* BL21(DE3) | *E. coli* strain used routinely for recombinant protein expression and purification. | Laboratory stock |

| | | |
|---|---|---|
| *M. smegmatis* mc$^2$155 | High-efficiency transformation strain of *M. smegmatis.* | Laboratory stock |
| pMV262~*mCherry* mc$^2$155 strain | *M. smegmatis* mc$^2$155 transformed with pMV262~*mCherry* replicative plasmid and is resistant to kanamycin. | This study |
| $P_{dnaE2}$~*mCherry* mc$^2$155 reporter | *M. smegmatis* mc$^2$155 transformed with 150bp upstream cloned of *M. tuberculosis dnaE2* gene containing its "SOS" box region upstream of *mCherry* reporter plasmid and is resistant to kanamycin. | This study |
| *M. tuberculosis* H37Ra | Avirulent, high-efficiency transformation strain of *M. tuberculosis.* | Laboratory stock |
| *M. tuberculosis* H37Rv | Virulent, high-efficiency transformation strain of *M. tuberculosis.* | Laboratory stock |
| *S. aureus* ATCC 25923 | Isolate used as a standard laboratory testing control strain. | Laboratory stock |
| **Plasmids/constructs**<br>pET28a (+) | Expression vector (pBR322 ori), strong phage promoter (T7), IPTG induction (lac operon), and kanamycin selection (kan$^r$). | Novagen |
| pMV262~*mCherry* | Mycobacterial replicative plasmid with pAL5000 mycobacterial origin of replication. *mCherry* is constitutively expressed from *hsp* promoter which is cloned between BamHI and EcoRI sites. The plasmid confers resistance to kanamycin. | Kind gift from Dr. Krishna Kurthkoti (RGCB, Trivandrum) |
| $P_{dnaE2}$~*mCherry* | 150bp upstream of *M. tuberculosis dnaE2* gene containing its "SOS" box region cloned upstream of *mCherry* between BamHI and NotI sites in pMV262~*mCherry* to create a "SOS" reporter plasmid resistant to kanamycin. | This study |
| S160A Mtb LexA | Catalytic site mutant of Mtb LexA which was used as a backbone for generating S160A/K197A Mtb LexA. | Kind gift from late Dr. M. Vijayan [1] (IISc, Bengaluru) |

**Table S3: Oligonucleotide primers used in this study**

| Primer for | Forward Primer (FP) / Reverse Primer (RP) Sequences |
|---|---|
| K197A Mtb LexA and S160A/K197A Mtb LexA | FP- 5´ GGCCACCGTCgcGACGTTCAAACG 3´<br>RP- 5´ TCACCGTCGATCATGGCC 3´ |
| *dnaE2*_44mer | As mentioned in [2] |

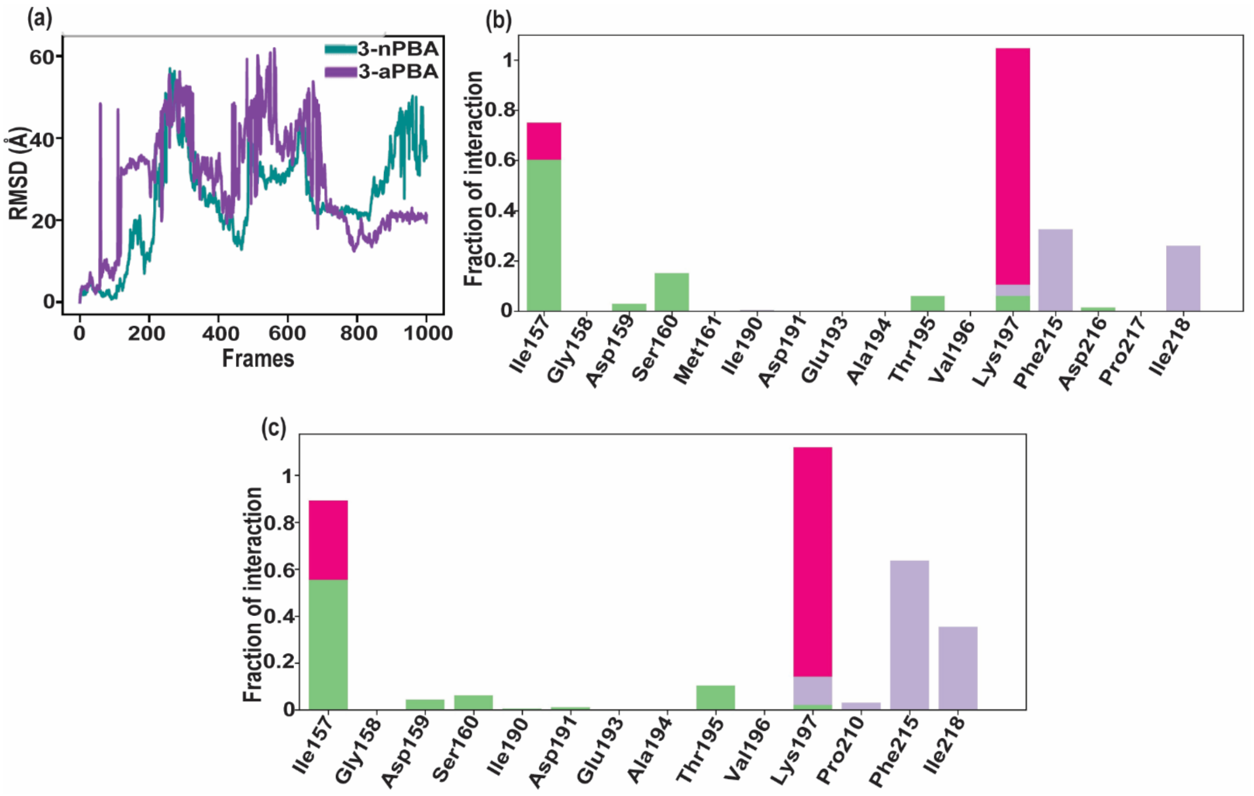


**Figure S1 a) Root Mean Square Deviation of 3-nPBA and 3-aPBA during 10 ns (1000 Frames) MD simulations of glide-docked complexes. b) Ligand atom interactions of 3-nPBA with Mtb LexA during 10ns run of covalently docked complex. shown as stacked bar charts. (c) Ligand atom interactions of 3-aPBA with Mtb LexA during 10ns run of covalently docked complex. shown as stacked bar charts. Hydrogen bonds are shown in green color, hydrophobic interactions in purple and ionic interactions in magenta color.**

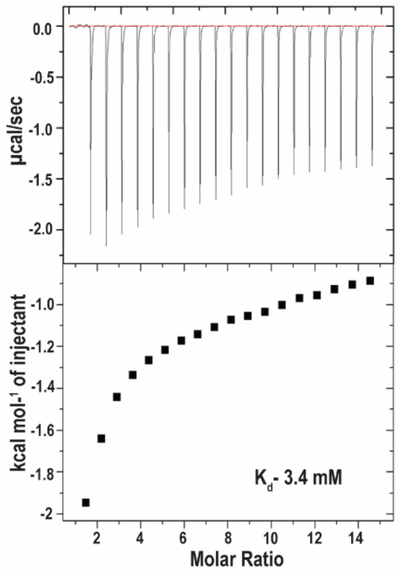


**Figure S2. Binding isotherm of S160A/K197A Mtb LexA with 3-nPBA.**

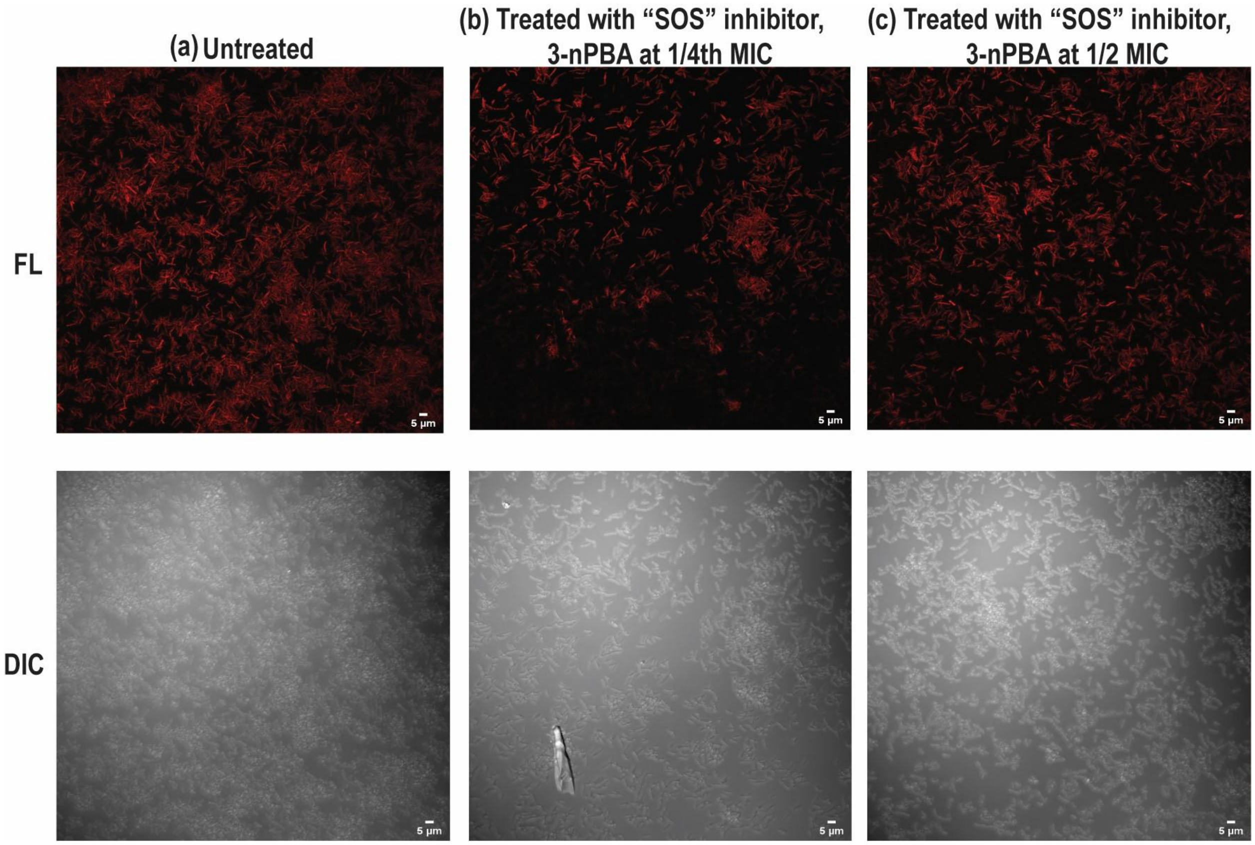


**Figure S3. Treatment with "SOS" inhibitor does not affect viability of cells. *M. smegmatis* cells transformed with pMV262~*mCherry* whereby *mCherry* gets constitutively expressed was used for this study. Treatment with the "SOS" inhibitor at both (b) 1/4th and (c) 1/2 of its minimum inhibitory concentration (MIC) is not seen to affect cell viability when compared to the (a) untreated control as the cells continue to exhibit fluorescence, which is an indication of their viable state.**

### MIC Determination using Resazurin Reduction Assay

For this, standard procedure was followed for *M. smegmatis* [3]. After setting up the plates, they were incubated at 37ºC for 40 hours at 100 rpm post which, 0.2 mg/ml final concentration of resazurin was added with further incubation for 6 hours after which images were taken. For testing in Mtb H37Ra, the following protocol was followed. Briefly, 200 µl MilliQ was added to the perimeter wells of a 96-well plate. Cultures of Mtb H37Ra were grown up to $O.D_{600}$ 0.6 in replicates and were diluted to 0.02. 100 µl of growth media was added in the required wells. Drugs were added from individual stocks to achieve the required final concentrations in the wells. 100 µl of culture was added to the wells to make a total reaction volume of 200 µl. Incubation was done at 37°C for 7 days. 30 µl of 0.01 % resazurin was added (final concentration 0.0015 %), incubated for 24 hours, and imaged. For testing in *S. aureus*, replicates of cultures were grown up to

O.D$_{600}$ 0.6 and diluted to 0.0008. 80 µl of culture was added with 20 µl of drugs from a 5 X higher stock. Plates were incubated for 24 hours at 37°C without shaking. 30 µl of resazurin was finally added to a concentration of 0.02 % and incubated for 2 hours at 37°C without shaking, post which images were taken. The protocol remained the same when followed for *E. coli* BL21DE3, with the only change being that the plates were incubated for 16 hours instead of for 24 hours as done for *S. aureus.* In all cases, ciprofloxacin was used as a positive control. Other relevant controls were taken as required. All experiments were performed in biological replicates as well as in technical replicates.

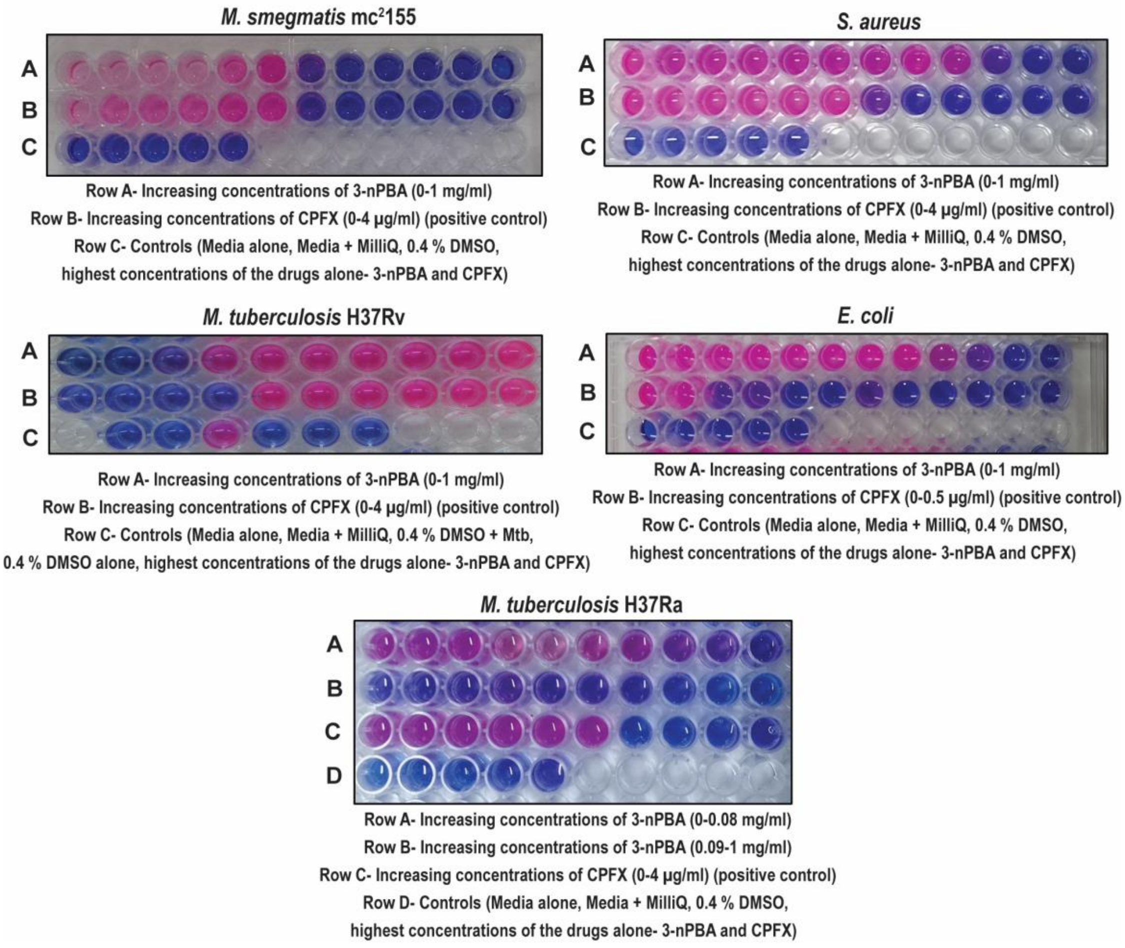


**Figure S4. Determination of MIC of "SOS" inhibitors (3-aPBA and 3-nPBA) and ciprofloxacin ("SOS" inducer) against mycobacterial species and representative Gram-positive and Gram-negative organisms.**

**Table S4: MICs against "SOS" inhibitors and "SOS" inducer**

| Molecules tested | 3-nPBA (μg/ml) | 3-aPBA (μg/ml) | Ciprofloxacin (μg/ml) |
|---|---|---|---|
| **MIC against *M. smegmatis*** | **30** | **625** | **0.125** |
| **MIC against Mtb H37Ra** | **90** | **1250** | **0.5** |
| **MIC against Mtb H37Rv** | **250** | **>2000** | **0.5** |
| **MIC against *E. coli* BL21 DE3** | **250** | **>2000** | **0.01** |
| **MIC against *S. aureus*** | **250** | **>2000** | **0.125** |

### Cytotoxicity assessment

The percentage of cell viability was assessed with Resazurin Reduction Assay in RAW 264.7 cell line by treating with the inhibitor. 10,000 cells/well were plated in a 96-well plate. After 24 h of cell plating, varying concentrations of 3-nPBA were applied. Treated cells were allowed to grow for another 48 h. Then, resazurin was added to each well (30 μl from a 0.01 % stock for a 200 μl reaction) and incubated for another 24 hours before taking readings. The percentage of cell viability was calculated.

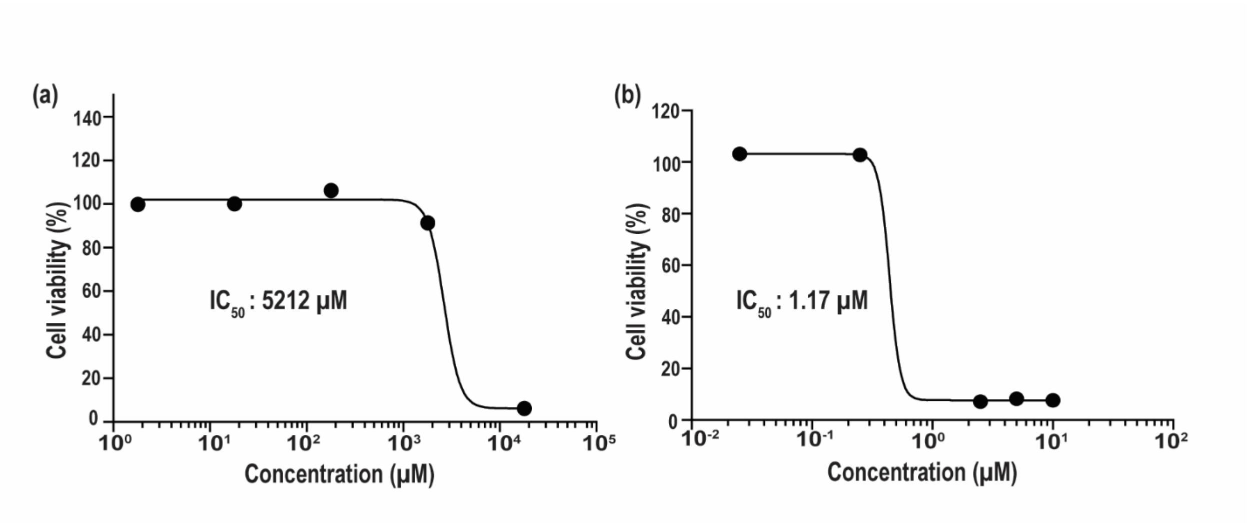


**Figure S5. Treatment with "SOS" inhibitor does not affect viability of RAW 264.7 cells (a). Doxorubicin was taken as positive control (b).**
